# Propagation, inactivation, and safety testing of SARS-CoV-2

**DOI:** 10.1101/2020.05.13.094482

**Authors:** Alexander S. Jureka, Jesus A. Silvas, Christopher F. Basler

## Abstract

In late 2019, a novel coronavirus, severe acute respiratory syndrome coronavirus 2 (SARS-CoV-2) emerged in Wuhan, the capital of the Chinese province Hubei. Since then, SARS-CoV-2 has been responsible for a worldwide pandemic resulting in over 4 million infections and over 250,000 deaths. The pandemic has instigated widespread research related to SARS-CoV-2 and the disease that it causes, COVID-19. Research into this new virus will be facilitated by the availability of clearly described and effective procedures that enable the propagation and quantification of infectious virus. Because work with the virus is recommended to be performed at biosafety level 3, validated methods to effectively inactivate the virus to enable safe study of RNA, DNA and protein from infected cells are also needed. Here, we report methods used to grow SARS-CoV-2 in multiple cell lines and to measure virus infectivity by plaque assay using either agarose or microcrystalline cellulose as an overlay as well as a SARS-CoV-2 specific focus forming assay. We also demonstrate effective inactivation by TRIzol, 10% neutral buffered formalin, beta propiolactone, and heat.

## 1. Introduction

The novel coronavirus SARS-CoV-2, the causative agent of Coronavirus disease 2019 (CoVID-19), belongs to the betacoronavirus genus, which also includes the highly pathogenic SARS CoV and MERS CoV. SARS-CoV-2, a close relative of bat betacoronaviruses emerged at the end of 2019 in Wuhan, China, and has caused a pandemic [1]. The pandemic and the public response has greatly impacted public health and caused profound social and economic disruption. As of May 11, 2020, SARS-CoV-2 had caused more than 4 million infections and more than 250,000 deaths worldwide [2]. The emergence of this new virus has prompted urgent worldwide efforts to develop diagnostics, vaccines and antivirals, to define the natural history of human infection; and to better characterize the virus.

The expanded interest in studying SARS-CoV-2 to address the current pandemic requires that many laboratories acquire the capacity to work with the virus. However, despite the rapidly growing body of literature either deposited on preprint servers or in peer-reviewed scientific journals, and the need for high quality data, there remains a lack of information regarding standardized protocols for work with the virus. Among these needs are the means to grow and quantify infectious virus. Additionally, the recommendation that experiments involving the propagation of SARs-CoV-2 be performed at biosafety level 3 (BSL3) necessitates the development of methods to safely inactivate the virus and validate inactivation methods for emerging highly-pathogenic viruses, such as SARS-CoV-2, to allow an array of studies to be performed at lower biocontainment levels [3-6]. Examples include the isolation of RNA from virus and virus-infected cells to characterize viral genome sequences, monitor viral gene expression and genome replication, and to characterize host responses to infection. Removal of intact, virus-infected cells is critical for studies involving microscopy aimed at understanding the SARS-CoV-2:host interplay at the cellular level, and also for high-throughput analysis of the progression of viral replication in response to antivirals. Whole inactivated virus and viral proteins are needed for the development of inactivated whole-virus vaccine preparations and also as a source of antigen for immunoassays.

To help address these needs and to facilitate SARS-CoV-2 research efforts, we describe here methods for the propagation of SARS-CoV-2 in multiple cell lines. We have also determined a more efficient method for quantifying virus by plaque assay and have developed a SARS-CoV-2-specific focus forming assay which can enhance throughput of assays requiring quantification of viral titers. Additionally, we describe validation if methods for the inactivation of SARS-CoV-2 through the use of TRIzol, 10% neutral buffered formalin, beta-propiolactone, and heat. Taken together, the data presented here will serve to provide researchers with a helpful basis of information to aid in their work on SARS-CoV-2.

## 2. Materials and Methods

### 2.1 Cells and Virus

Vero E6 (ATCC# CRL-1586), Calu-3 (ATCC# HTB-55), Caco-2 (ATCC# HTB-37), Huh7, A549 (ATCC# CCL-185), and 293T cells were maintained in DMEM (Corning) supplemented with 10% heat inactivated fetal bovine serum (FBS; GIBCO). Cells were kept in a 37°C, 5% CO_2_ incubator without antibiotics or antimycotics. SARS-CoV-2, strain USA_WA1/2020, was obtained from the World Reference Collection for Emerging Viruses and Arboviruses at the University of Texas Medical Branch-Galveston.

### 2.2 Virus Propagation

A lyophilized ampule of SARS-CoV-2 was initially resuspended in DMEM supplemented with 2% FBS. VeroE6 cells were inoculated in duplicate with a dilution of 1:100 with an adsorption period of 1 hour at 37°C and shaking every 15 minutes. Cells were observed for cytopathic effect (CPE) every 24 hours. Stock SARS-CoV-2 virus was harvested at 72 hours post infection (h.p.i) and supernatants were collected, clarified, aliquoted, and stored at −80°C. For replication kinetic experiments, cells were seeded into 24 well plates at confluency in DMEM supplemented in 10% FBS. The next day, cells were inoculated with SARS-CoV-2 at a multiplicity of infection (MOI) of 0.01 in DMEM supplemented with 2% FBS. Supernatants were harvested at the indicated timepoints and stored at −80C until analysis.

### 2.3 SARS-CoV-2 Plaque Assay

Vero E6 cells were seeded into 6, 12, or 24-well plates 24 hours before infection. 10-fold serial dilutions of SARS-CoV-2 samples were added, adsorbed for 1 hour at 37C with shaking at 15-minute intervals. After the absorption period, 2 mL of 0.3% agarose in DMEM supplemented with 2% FBS or 3 mL of 0.6 or 1.2 percent microcrystalline cellulose (MCC; Sigma 435244) in serum-free DMEM was added. To stain plaque assays performed with an agarose overlay, 10% Neutral Buffered Formalin (NBF) was added on top of the agarose and incubated for one hour at room temperature. The agarose plug was then removed with a pipette tip, and the fixed monolayer was stained with 0.4% crystal violet in 20% methanol. For plaque assays performed with MCC overlay, the MCC was aspirated out, 10% NBF added for one hour at room temp and then removed. Monolayers were then washed with water and stained with 0.4% crystal violet. Plaques were quantified and recorded as PFU/mL.

### 2.4 Focus Forming Assay

Vero E6 cells were plated into 96 well plates at confluency (75000 cells/well) in DMEM supplemented with 10% heat-inactivated fetal bovine serum (Gibco). Prior to infection, virus stocks were thawed and serially diluted to obtain dilutions in the range of 10^−2^ to 10^−9^. Growth media was removed from the Vero E6 cells and 50 µL of virus dilutions was plated. Virus was adsorbed for 1 hour at 37C/5% CO2. After adsorption, 50 µL of 2.4% MCC overlay supplemented with DMEM powdered media to a concentration of 1X (Gibco) was added to each well of the 96 well plate to achieve a final MCC overlay concentration of 1.2%. Plates were then incubated at 37C/5% CO2 for 24 hours. The MCC overlay was gently removed and cells were fixed with 10% NBF for 1 hour at room-temperature. After removal of NBF, monolayers were washed with ultrapure water and 100% methanol/0.3% H_2_O_2_ was added to permeabilize cells and quench endogenous peroxidase activity. Monolayers were then blocked for 1 hour in PBS with 5% non-fat dry milk (NFDM). After blocking, monolayers were incubated with SARS-CoV N primary antibody (Novus Biologicals; NB100-56576 – 1:2000) for 1 hour at RT in PBS/5% NFDM. Monolayers were washed with PBS and incubated with an HRP-Conjugated secondary antibody for 1 hour at RT in PBS/5% NFDM. Secondary was removed, monolayers were washed with PBS, and then developed using TrueBlue substrate (KPL) for 30 minutes. Plates were imaged on a Bio-Rad Chemidoc utilizing a phosphorscreen and foci were quantified.

### 2.5 TRIzol® Treatment

To validate the effectiveness of Trizol® Reagent in the inactivation of SARS-CoV-2, stock virus was separated into the following validation test groups: non-infected control (sterile media), Trizol treated non-infected control (sterile culture media+Trizol), duplicate SARS-CoV-2 positive controls (SARS-CoV-2; 1×10^6^ pfu), and triplicate SARS-CoV-2 Trizol treated samples (SARS-CoV-2; 1×10^6^ pfu +Trizol). Trizol was added to a final concentration of 10%. All test groups were then incubated at room temperature for 10 minutes. Samples were then diluted in 25mL media, and plated onto VeroE6 cells in a 15cm tissue culture dish and incubated at 37C and 5% CO2 and monitored daily for CPE. At 48 hours post infection, media was removed and the plates fixed with 10% NBF. To determine integrity of the cell monolayer, plates were stained with crystal violet.

### 2.6 Formalin Treatment

To validate inactivation of SARS-CoV-2 with formalin, 21 wells of a 24-well plate seeded to confluency with VeroE6 cells were infected with SARS-CoV-2 at an MOI of 0.1. Three wells served as mock-infected control. At 24 hours post infection, media was removed from wells and treated as follows, using 3 wells per condition: uninfected control, SARS-CoV-2 infected control, SARS-CoV-2+2% formaldehyde, SARS-CoV-2+1% formaldehyde, SARS-CoV-2+0.5% formaldehyde, SARS-CoV-2+0.1% formaldehyde, and SARS-CoV-2+0.05% formaldehyde. Formaldehyde treatment was carried out at room temperature for 1 hour. The cells were then washed with fresh media, scraped off the plate, and overlaid onto uninfected Vero E6 cells. Samples were then incubated at 37C, 5% CO2 and monitored daily for CPE. At 72 hours post infection, media was removed and the plates fixed with 10% NBF. To determine integrity of the cell monolayer, plates were stained with crystal violet.

### 2.7 Beta-propiolactone Treatment

To test treatment of viral particles with beta-propiolactone as means of inactivation, stock SARS-CoV-2 virus was separated into the following validation groups: non-infected control, 4C control (1×10^6^ pfu per replicate), 0.5% beta-propiolactone (1×10^6^ pfu per replicate), 0.1% beta-propiolactone (1×10^6^ pfu per replicate), and 0.05% beta-propiolactone (1×10^6^ pfu per replicate). After incubation at 4C for 16 hours, samples were transferred to 37C for two (2) hours to hydrolyze all residual beta-propiolactone. This step ensures complete hydrolysis of beta-propiolactone to prevent cytotoxicity to mammalian cells [7]. After hydrolysis, samples were inoculated onto Vero E6 cells, and incubated at 37C, 5% CO2 incubator and monitored daily for CPE. At 72 hours post infection, media was removed and plates were fixed with 10% NBF. To determine integrity of the cell monolayer, plates were stained with crystal violet.

### 2.8 Heat Inactivation

To validate heat treatment as method to inactivate SARS-CoV-2, SARS-CoV-2 virus was separated into the following validation groups: non-infected control, room temperature control (1×10^6^ pfu per replicate), 100C for 5 minutes (1×10^6^ pfu per replicate), 100C for 10 minutes (1×10^6^ pfu per replicate), and 100C for 15 minutes (1×10^6^ pfu per replicate). 1.5 ml microcentrifuge polypropylene tubes containing the virus (500 µL total volume) were exposed to direct heat in a heat block (Fisher Scientific). After heating, all samples were left to cool to room temperature and centrifuged to collect condensation within the tube. Each sample was then inoculated in 5-fold serial dilutions onto VeroE6 cells. Samples were then incubated at 37C, 5% CO2. At 72 hours post infection, supernatants were harvested, and infectious virus was quantified by plaque assay on Vero E6 cells.

### 2.9 Electron Microscopy

Resuspended purified beta-propiolactone treated SARS-CoV-2 virus preps were adsorbed onto 300 mesh formvar-carbon coated nickel grids for 10 minutes, washed with 0.2 µM filtered ultrapure water, and negative-stained with UranyLess stain for 15 seconds. Grids were then washed with 0.2 µM ultrapure water, allowed to dry, and imaged on a LEO 960 TEM at 80 kV.

### 2.10 Western Blotting

Resuspended purified beta-propiolactone treated SARS-CoV-2 virus preps were separated by SDS-PAGE (Bio-Rad TGX Mini) and transferred to 0.2 µM PVDF membrane according to the manufacturer’s protocols (Bio-Rad Tansblot Turbo). After blocking in 5% non-fat dry milk in TBST (10 mM Tris, 150 mM NaCl, 0.5% Tween-20, pH8) for one hour, membranes were incubated overnight at 4C with antibodies targeting SARS-CoV N (Novus Biologicals; NB100-56576) or SARS-CoV S (Sino Biological; 40150-T62). Membranes were washed in TBST and incubated with an HRP-conjugated rabbit secondary antibody (Cell signaling; 7074) for 1 Hr at room temperature. Membranes were then washed, developed with ECL, and imaged on a Bio-Rad Chemidoc imaging system.

## 3. Results

### 3.1 Propagation and quantification of SARS-CoV-2 in cell culture

Despite the number of recent reports in which SARs-CoV-2 has been propagated and quantified, there still remains a lack of general information regarding the temporal cytopathology of SARS-CoV-2 in cell culture. Here, we set out to determine the most appropriate times post-infection to harvest SARS-CoV-2 infected cultures for generation of stock virus, and the quantification thereof. To generate stock virus, Vero E6 cells were infected with SARS-CoV-2 at an MOI of 0.001 and monitored daily by light microscopy for the appearance of CPE. We determined that the cytopathic effect (CPE) caused by SARS-CoV-2 in Vero E6 cells is most apparent at 48 hours post infection (Figure 1A). However, despite CPE being nearly complete at 48 hours post-infection in the absence of an overlay, we determined that for the quantification of SARS-CoV-2 using traditional plaque assays with a 0.4% agarose overlay 72 hours post-infection results in clearer, more easily quantifiable plaques (Figure 1B).

**Figure 1.**
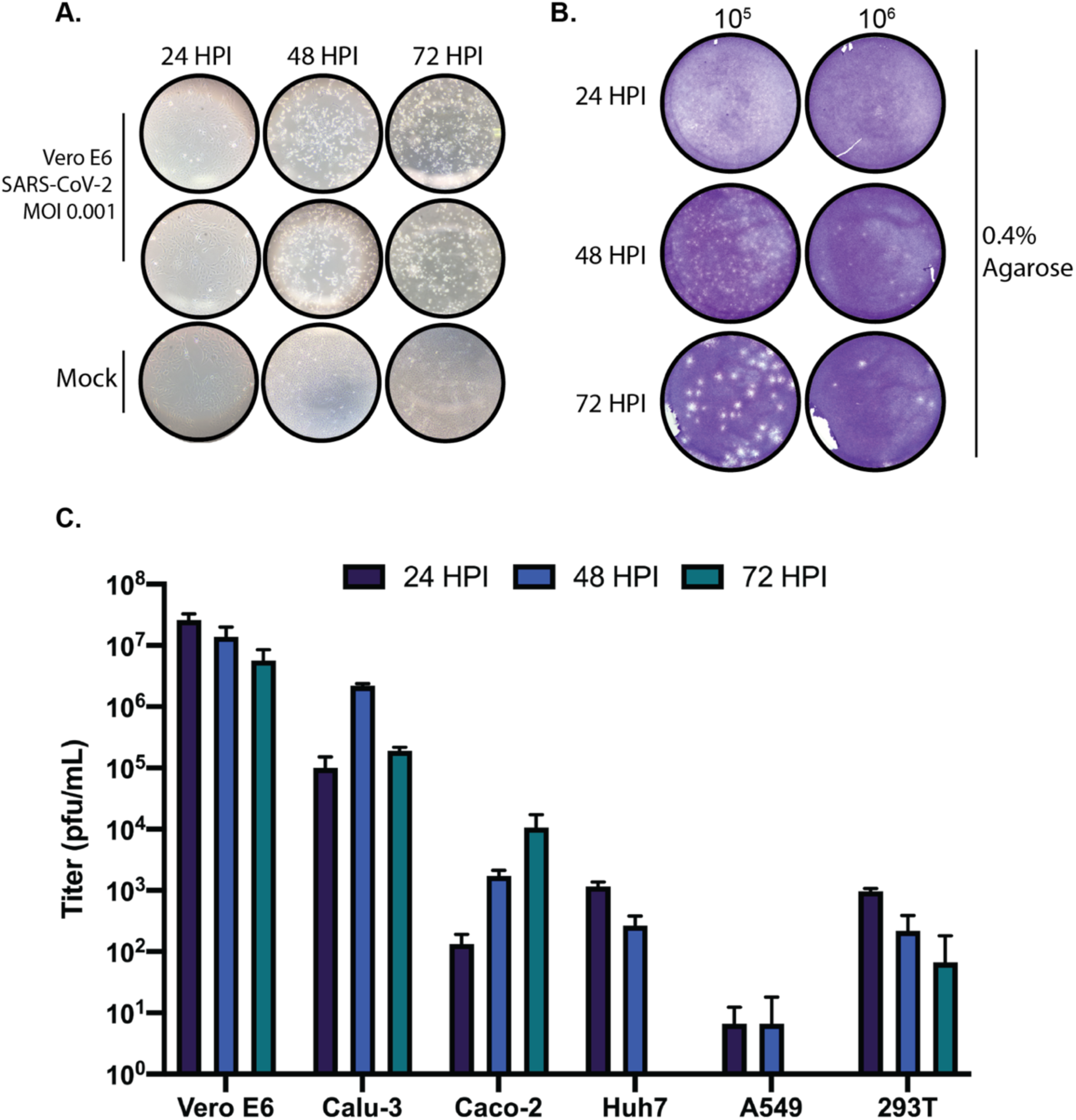
Propagation and quantification of SARS-CoV-2 in cell culture. (A) Vero E6 cells were infected with SARS-CoV-2 at an MOI of 0.001 and monitored daily by microscopy for the presence of CPE. (B) Vero E6 cells were infected with serially diluted SARS-CoV-2 stock virus and overlaid with 0.4% agarose in DMEM supplemented with 2% FBS. Plaque assays were harvested at 24, 48, and 72 hours post-infection, fixed, and stained with crystal violet to visualize plaques. (C) Vero E6, Calu-3, Caco-2, Huh7, A549, and 293T cells were infected with SARS-CoV-2 at an MOI of 0.01. Supernatants from 24, 48, and 72-hour timepoints were quantified by plaque assay. Data are representative of the mean and SEM of 3 replicates.

In addition, we wanted to determine the kinetics of SARS-CoV-2 in several commonly used cell lines to determine suitable cell culture systems for studying SARS-CoV-2 biology. Vero E6, Calu-3, Caco-2, Huh7, A549, and 293T cells were infected with SARS-CoV-2 at an MOI of 0.01 and supernatants were harvested at 24, 48, and 72 hours post-infection and quantified by traditional plaque assays. As shown in Figure 1C, SARS-CoV-2 replicated to high titers in Vero E6 and Calu3 cells at all times post-infection. Caco-2 cells clearly support SARS-CoV-2 replication yet seem to propagate the virus more slowly that Vero E6 or Calu-3 cells. Only modest replication was observed in Huh7 and 293T cells, and A549 cells did not support SARS-CoV-2 growth at any time post-infection. These results suggest that the cellular tropism for SARS-CoV-2 is fairly restricted; however overexpression of the SARS-CoV-2 receptor ACE2 or the protease responsible for cleaving SARS-CoV-2 spike protein can aid infection in non-permissive cell lines [8,9].

### 3.2 Microcrystalline cellulose is a suitable substitute for agarose for SARS-CoV-2 plaque assays

Traditionally, viral plaque assays are performed utilizing low concentrations of agarose as an overlay medium. While agarose generally performs well as an overlay medium for numerous different viruses, the requirement of having to remove agarose plug from within multi-well cell culture dishes can prove laborious. This is especially the case when plaque assays are performed in 24 well plates. Recently, a novel low-viscosity overlay medium for viral plaque assays has been described [10]. This assay replaces traditional solid (agarose) and semi-solid (methylcellulose) media with a microcrystalline cellulose (MCC) suspension as the overlay media. To determine whether the MCC approach is suitable for quantifying SARS-CoV-2 by plaque assay, we overlaid SARS-CoV-2 infected Vero E6 cells with two concentrations of MCC (0.6% and 1.2%) in 6, 12, and 24 well formats. Our results indicate that MCC performs exceptionally well as an overlay medium for SARS-CoV-2 plaque assays as uniform and countable plaques are readily apparent 72 hours post-infection, even in the 24 well plate format (Figure 2 A-B). We have also determined that either of the two concentrations of MCC tested yielded plaques of comparable size. Based on this we would recommend the use of 0.6% MCC overlay media (Figure 2A) for SARS-CoV-2 plaque assays as it produces uniform and easily identifiable plaques for quantification. While it is possible to identify plaques at 48 hours post-infection with 0.6% MCC overlay, similar to the 0.4% agarose overlay (Figure 1B), the plaques are small and difficult to accurately quantify until 72 hours post-infection. Taken together these data demonstrate that using MCC as an overlay media is an effective and far more efficient method than the use of traditional agarose overlays.

**Figure 2.**
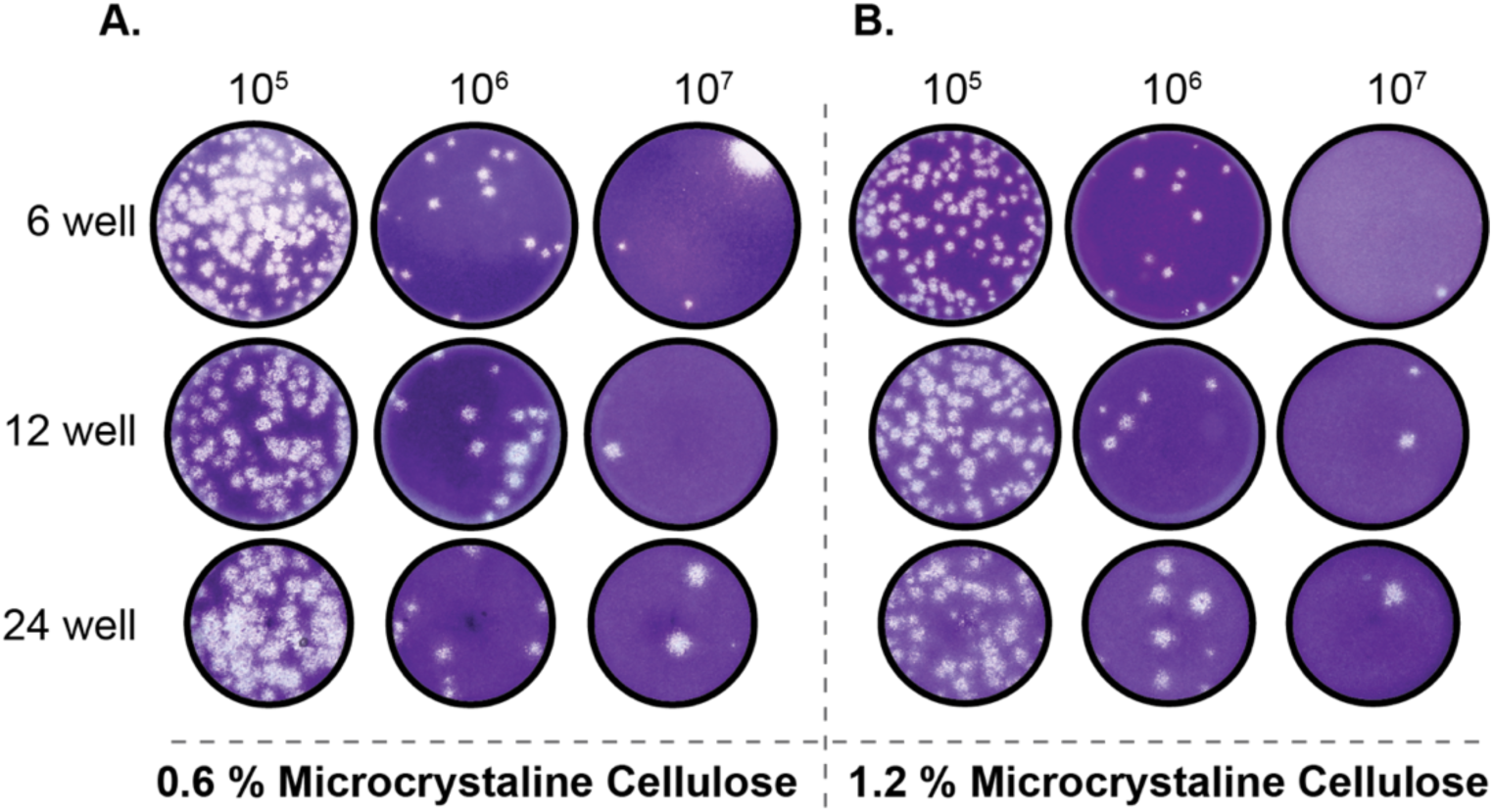
MCC is a suitable alternative as an overlay medium for SARS-CoV-2 plaque assays. Vero E6 cells were infected with serially diluted SARS-CoV-2 stock virus and overlaid with 0.6% (A) or 1.2% (B) MCC in serum free DMEM. After 72 hours, the MCC overlay was removed and monolayer were fixed and stained with crystal violet to visualize plaques.

### 3.3 Development of a foci forming assay for SARS-CoV-2

For viruses like SARS-CoV-2 which produce significant CPE in permissive cell lines, traditional plaque assays are the standard for virus quantification. However, traditional plaque assays require waiting for a particular virus to produce significant enough CPE for quantifiable plaque formation. As described above, SARS-CoV-2 is most readily quantifiable by plaque assay at 3 days post-infection. Here, we set out to develop and immunohistochemical assay to reliably determine SARS-Cov-2 titers using a 96 well plate-based foci forming assay. Our results indicate that after only 24 hours post infection, SARS-CoV-2 foci are readily detectable and quantifiable (Figure 3A). Although the titers obtained by the foci forming assay after 24h incubation are approximately 1 log lower than titers obtained from traditional plaque assays at 72h,, both data sets share similar trends (Figure 3B). Taken together, these data demonstrate that SARS-CoV-2 containing samples can be accurately quantified within 24 hours, and in a higher throughput manner than traditional plaque assays.

**Figure 3.**
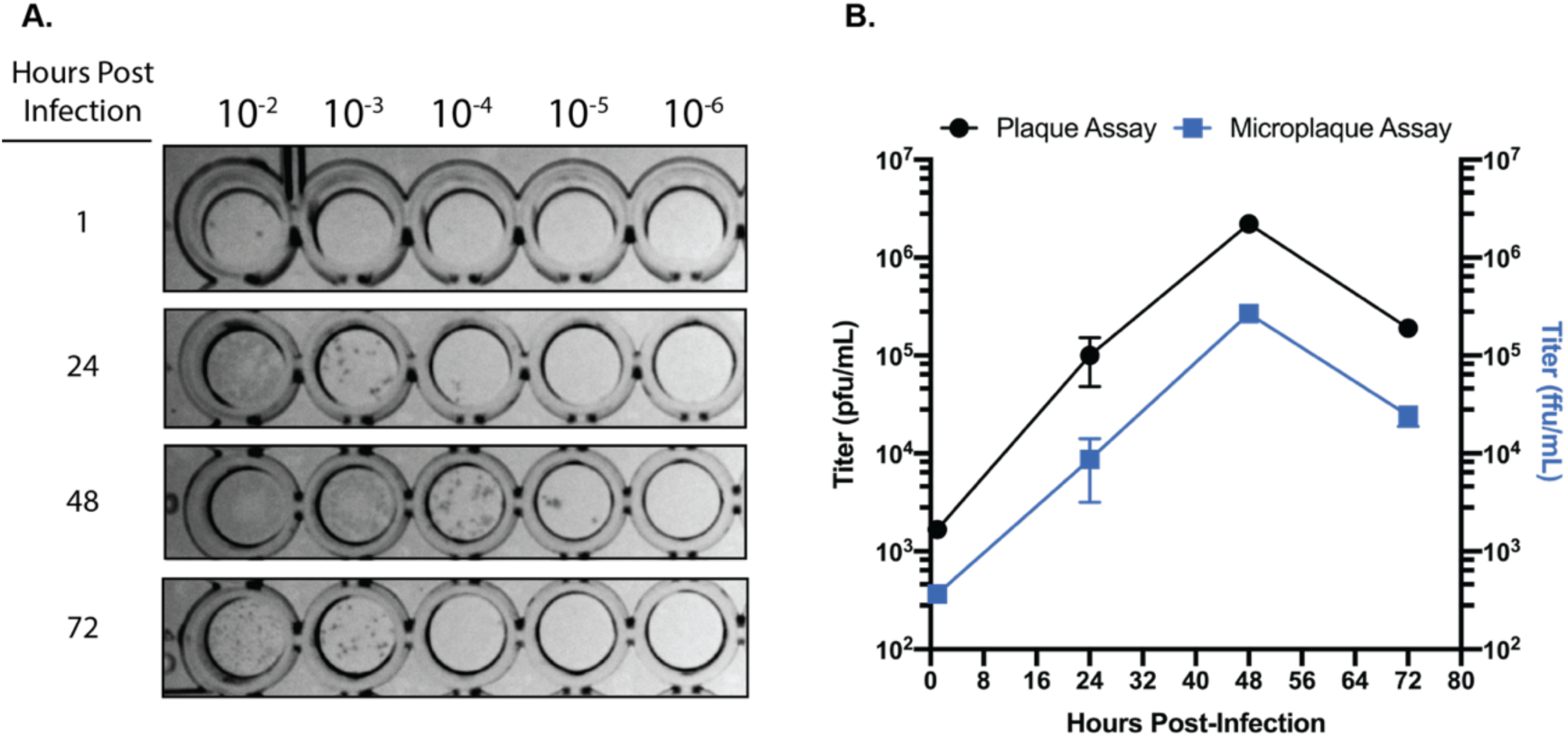
SARS-CoV-2 focus forming assay. (A) Representative image of the foci observed from serially diluted supernatants from infected Calu-3 cells. (B) Comparison of titers obtained from traditional plaque assay and focus forming assays on matched supernatants collected from SARS-CoV-2 infected Calu-3 cells. Data are representative of the mean and SEM of 3 replicates.

### 3.4 Validation of inactivation of SARS-CoV-2 by TRIzol, formalin, beta-propiolactone, and direct heat

Currently, there is limited data published on methods that successfully inactivate SARS-CoV-2. To help fill this gap in knowledge, we tested the ability of TRIzol, formalin, beta-propiolactone, and direct heat to successfully inactivate SARS-CoV-2.

#### 3.4.1 TRIzol

TRIzol is a well-known and widely used reagent for the isolation of nucleic acids and in some cases protein from cells. TRIzol treated samples are generally very stable, and the ability to isolate RNA/DNA/protein without the need for column-based extraction kits with proprietary task-specific lysis buffers makes it an attractive option for processing SARS-CoV-2 infected samples. We determined that addition to SARS-CoV-2 stock virus (1×10^6^ pfu; see Materials and Methods) of TRIzol to a final concentration of 10% resulted in the successful inactivation of SARS-CoV-2 as treated samples resulted in no detectable cytopathic effect on Vero E6 cells 72 hours post-infection (Table 1).

**Table 1.** Inactivation of SARS-CoV-2 by TRIzol^®^, formaldehyde, and beta-propiolactone

| Treatment |  | CPE (72 Hours Post-Infection) |  |  |
| --- | --- | --- | --- | --- |
| Trizol® | Input (pfu) | Replicate 1 | Replicate 2 | Replicate 3 |
| 10% | 1x10<sup>6</sup> | ND | ND | ND |
| **Formaldehyde** |  |  |  |  |
| 2% | Infected Cells<br>(MOI 0.1) | ND | ND | ND |
| 1% | Infected Cells<br>(MOI 0.1) | ND | ND | ND |
| 0.5% | Infected Cells<br>(MOI 0.1) | ND | ND | ND |
| 0.1% | Infected Cells<br>(MOI 0.1) | CPE | ND | ND |
| 0.05% | Infected Cells<br>(MOI 0.1) | CPE | CPE | CPE |
| **Beta-propiolactone** |  |  |  |  |
| 0.5% | 1x10<sup>6</sup> | CD | CD | CD |
| 0.1% | 1x10<sup>6</sup> | CD | ND | ND |
| 0.05% | 1x10<sup>6</sup> | ND | ND | ND |
| **CPE present (CPE), CPE not detected (ND), Cell death (CD)** |  |  |  |  |

#### 3.4.2 Formalin

10% neutral buffered formalin is a commonly used fixative for processing infected cells for downstream analysis such as microscopy. In order to determine the effectiveness of formaldehyde in inactivating SARS-CoV-2, Vero E6 cells were infected at an MOI of 0.01 and incubated at 37C for 24 hours. After incubation, media was removed, and cells were treated with varying concentrations of formaldehyde for 1 hour at room temperature. Fixed cells were then scraped off the dish and transferred to freshly plated Vero E6 cells. Our results indicate that formalin is effective in inactivating SARS-CoV-2 at concentrations ranging from 0.5% to 2% (total formaldehyde concentration) after 1 hour at room temperature (Table 1). While the exact viral load inactivated was not determined, the impressive viral titers observed on Vero E6 cells at 24 hours post-infection (Figure 1C) suggests the inactivation described here shared a similar viral burden to the others in this report. Given that 10% neutral buffered formalin (NBF) contains 4% formaldehyde, treatment of SARS-CoV-2 containing samples with 10% NBF for 1 hour at room temperature is more than adequate for SARS-CoV-2 inactivation. Our data also indicates that lower concentrations (down to 0.5% total formaldehyde concentration) will effectively inactivate SARS-CoV-2 for the purposes of processing of liquid samples, such as isolation of whole virions.

#### 3.4.3 Beta-propiolactone

Beta-propiolactone (BPL) is a commonly used reagent for the inactivation of viruses for use in vaccine preparations [11-14] and it has recently been used in the development of an inactivated SARS-CoV-2 vaccine preparation [15]. Our results indicate that incubation of SARS-CoV-2 (×10^6^ pfu) in solution with 0.5% BPL for 16 hours at 4C followed by a 2-hour incubation at 37C results in complete inactivation of infectious SARS-CoV-2 (Table 1). Examination of control cells treated with 0.5% BPL by light microscopy and crystal violet staining indicated no apparent cytotoxicity 72 hours post-treatment.

Given its use in the preparation of vaccines, we wanted to determine if BPL could provide a rapid method for the purification of inactivated viral particles. Our results indicate that after purification of BPL treated SARS-CoV-2 stocks over a 20% sucrose gradient (Figure 4A) that intact viral particles are readily apparent by electron microscopy (Figure 4B). We also identified that both nucleoprotein and spike of SARS-CoV-2 are detectable by western blot in these samples. These results indicate that BPL inactivation of SARS-CoV-2 viral particles and their subsequent purification will yield inactivated, intact viral particles.

**Figure 4.**
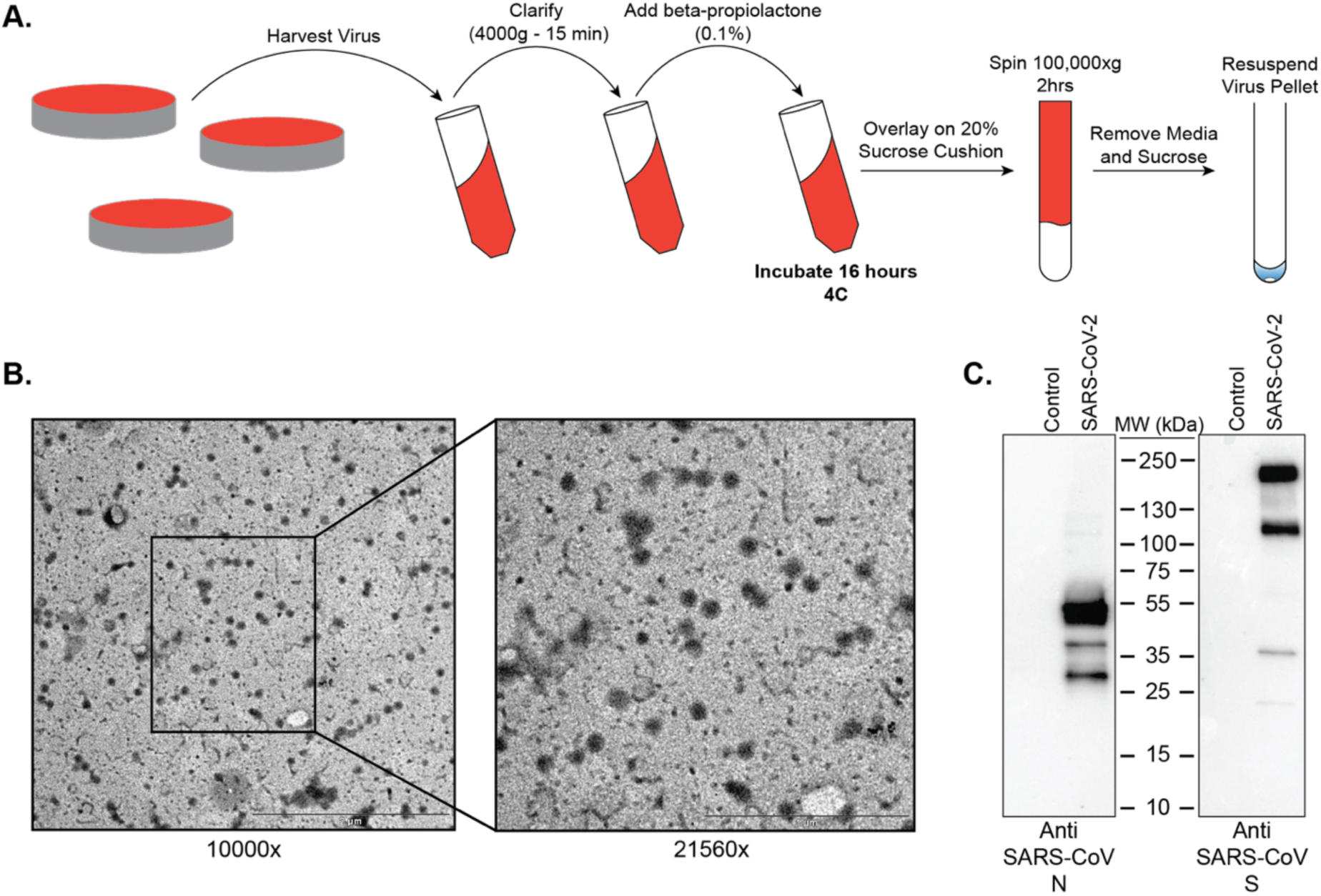
BPL inactivation and purification of SARS-CoV-2 virus particles. (A) Graphical depiction of the workflow established for the inactivation and purification of BPL inactivated SARS-CoV-2. (B) Negative stain TEM images of SARS-CoV-2 virus particles after purification from cell culture media over a 20% sucrose cushion at 100,000 xg for 2 hours. (C) Western blots of BPL inactivated virus particles for SARS-CoV-2 nucleoprotein and spike protein.

#### 3.4.4 Heat

To determine the effectiveness of heat in inactivating SARS-CoV-2 with respect to time, we heated virus stocks at 100C for 5, 10, and 15 minutes and 56C for 15, 30, 45, and 60 minutes. After heating, remaining infectious SARS-CoV-2 was quantified by plaque assay. Treatment of SARS-CoV-2 (1×10^6^ pfu) for 5 minutes at 100C and 45 minutes at 56C resulted in complete inactivation of infectious virus (Figure 5).

**Figure 5.**
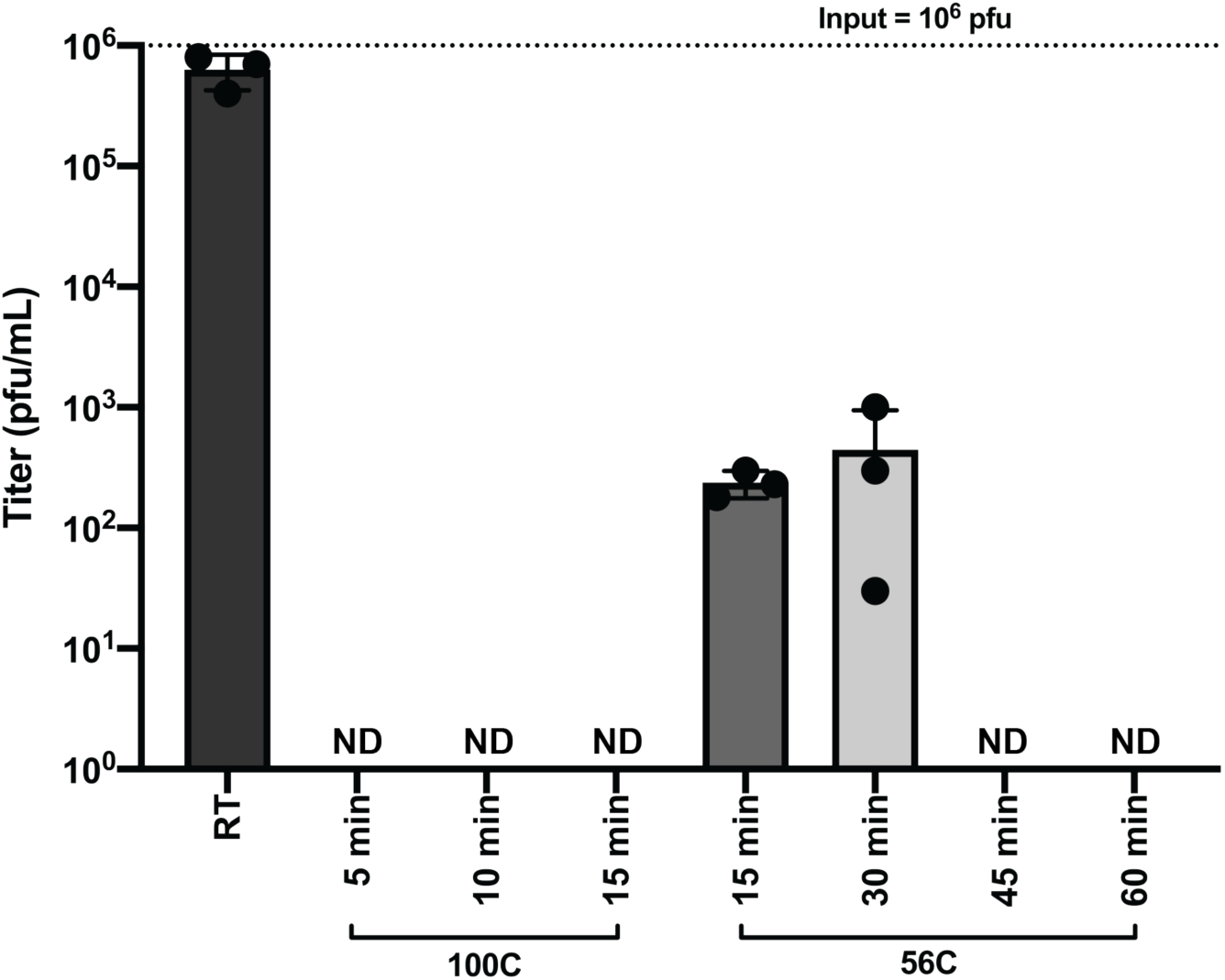
Heat Inactivation of SARS-CoV-2. SARS-CoV-2 containing samples (1×10^6^ pfu) were heated at 100C for 5, 10 and 15 minutes and 56C for 15, 30, 45, and 60 minutes. Samples were assayed by plaque assay to detect remaining infectious virus post-heating. The room temperature control was incubated at room temperature until all heated samples were prepared. Data are representative of the mean and SEM of 3 replicates.

## 4. Discussion

The ability to grow and accurately quantify infectious virus is critical for virological studies. Here, we sought to determine the growth kinetics of SARS-CoV-2 in several commonly used cell lines as well as the most appropriate time post-infection to accurately quantify SARS-CoV-2 by plaque assay. Significant growth was achieved in both Vero E6 and Calu-3 cells at the time points tested. We also observed that the colorectal adenocarcinoma Caco-2 cell line was able to propagate SARS-CoV-2, albeit to lower titers than Vero E6 and Calu-3 cells. Based on the replication kinetics observed in Caco-2 cells, it is likely that if the infection were allowed to progress past 72 hours that higher titers would have been achieved. Huh7 and 293T cells exhibited modest growth of SARS-CoV-2, and A549 cells did not support SARS-CoV-2 replication at any of the tested timepoints. Therefore, these cell lines are not ideal hosts for virological studies with SARS-CoV-2 in the absence of modifications such as overexpression of the SARS-CoV-2 entry receptor ACE2 or the cellular protease TMPRSS2 which is necessary for Spike processing [8,9].

Plaque assays are among the most commonly used techniques for the accurate quantification of infectious virus. To establish reliable plaque assays for SARS-CoV-2 we compared two separate approaches. One used an agarose overlay and the other used microcrystalline cellulose (MCC). In our hands, both approaches yielded comparable data. However, the MCC overlay approach has significant advantages over the more traditional agarose overlay. MCC overlays do not require heating prior to use which significantly expedites the processing of large numbers of plaque assays and also alleviates concerns of overheating cell monolayers. Additionally, MCC overlays do not require the removal of a “plug” like in the use of agarose overlays as MCC overlays remain liquid throughout their use. They can simply be aspirated, and monolayer can be further processed. However, due to the fact that MCC overlays remain liquid throughout the course of the assay, it is important to take care that plaque assays utilizing MCC as an overlay are not moved until harvested to ensure reliable data. Our data presented here demonstrate that MCC is an effective and efficient overlay alternative to agarose for SARS-CoV-2 plaque assays.

While plaque assays are perhaps the most commonly used technique for the quantification of infectious virus, immunohistochemical focus forming assays are also commonly used for the quantification of virus [10,16-18]. We have established that SARS-CoV-2 can be reliably quantified by a 96 well plate-based focus forming assay in only 24 hours. When compared to traditional plaque assays for SARS-CoV-2 which require 72 hours for easily quantifiable plaques, the ability to quantify infectious SARS-CoV-2 within 24 hours in a 96 well plate format represents a significant advantage for studies requiring higher-throughput. While it is important to acknowledge that titers acquired with a focus forming assay will generally be one log lower than those acquired by traditional plaque assay, the focus forming assay described here is fully capable of differentiating log-scale changes in virus titer. Taken together, the focus forming assay described here is a rapid and efficient method for quantifying infectious SARS-CoV-2.

Currently the recommendation from the Centers for Disease Control and Prevention (CDC) is for studies involving the propagation of SARS-CoV-2 to be performed within BSL3 laboratories with standard BSL3 practices. While an official risk group determination has not been made for SARS-CoV-2, the related viruses SARS-CoV and MERS are classified as risk-group 3 pathogens. As a BSL3 pathogen, having validated methods to inactivate SARS-CoV-2 is important so as to ensure safety while also allowing removal of samples to lower biosafety levels for analysis. For example, whole inactivated virus can serve as antigen for immunological assays, vaccine preparations, and for the analysis of virus composition. We chose to examine beta-propiolactone as a means to inactivate SARS-CoV-2. This method was chosen because it preserves virus structure and antigenicity, and it has recently been used to generate an inactivated vaccine preparation for SARS-CoV-2 [15]. Our data show that treatment of SARS-CoV-2 with beta-propiolactone at a concentration of 0.5% for 16 hours at 4C followed by 2 hours at 37C will yield intact viral particles that can be utilized safely for downstream purposes.

The isolation of viral RNA and protein from virus or virus infected cells or the isolation RNA from host cells is critical to characterize virus sequence variation and to study the impact of infection on host gene expression. TRIzol is a commonly used reagent for RNA isolation that can also be used to for DNA and protein isolation. Our data show that treatment of SARS-CoV-2 containing samples with TRIzol following the manufacturer’s instructions is an effective method of inactivating SARS-CoV-2.

Formaldehyde is a ubiquitously used reagent in life sciences and has numerous applications such as the fixation of cells for microscopy studies such as indirect immunofluorescence of infected cells and as a disinfectant for scientific equipment. Here, we demonstrate that treatment of SARS-CoV-2 infected cells with formaldehyde at a concentration at or above 0.5% for one hour at room temperature effectively inactivates SARs-CoV-2. Given the wide ranges of uses for formaldehyde, we hope this data will assist in the safe processing of probable or confirmed SARS-CoV-2 containing samples.

Analyses of host responses to viral infection either through the use of Western blotting or the determination of virus specific antibodies in patient sera generally require heating of the sample prior to downstream analysis. To this end, we have determined that heating of SARS-CoV-2 containing samples at 100C for longer than 5 minutes is a safe and effective method for the downstream analysis of samples by western blot. Likewise, our data demonstrate that heat treatment of laboratory or clinical samples at 56C for one hour will serve to effectively inactivate SARS-CoV-2.

Taken together, we hope the methods and data reported here will serve to expedite the much-needed research required to address this unprecedented pandemic.

## Funding

This research was funded by NIH grants to CFB, including P01AI120943 and R01AI143292.

### Acknowledgments

We would like to thank Natasha K. Griffith, Marty Wildes, and Michael Walsh of the GSU High Containment Core for their support in completing the studies reported here.

## References

1. Bedford, J.; Enria, D.; Giesecke, J.; Heymann, D.L.; Ihekweazu, C.; Kobinger, G.; Lane, H.C.; Memish, Z.; Oh, M.D.; Sall, A.A., et al. COVID-19: towards controlling of a pandemic. Lancet 2020, 395, 1015–1018, doi: 10.1016/S0140-6736(20)30673-5.

2. Situation report - 75 Coronavirus disease 2019 (COVID-19) 4 April 2020. World Health Organization 2020.

3. Blow, J.A.; Dohm, D.J.; Negley, D.L.; Mores, C.N. Virus inactivation by nucleic acid extraction reagents. J Virol Methods 2004, 119, 195–198, doi: 10.1016/j.jviromet.2004.03.015.

4. Darnell, M.E.; Subbarao, K.; Feinstone, S.M.; Taylor, D.R. Inactivation of the coronavirus that induces severe acute respiratory syndrome, SARS-CoV. J Virol Methods 2004, 121, 85–91, doi: 10.1016/j.jviromet.2004.06.006.

5. Haddock, E.; Feldmann, F.; Feldmann, H. Effective Chemical Inactivation of Ebola Virus. Emerg Infect Dis 2016, 22, 1292–1294, doi: 10.3201/eid2207.160233.

6. Rabenau, H.F.; Cinatl, J.; Morgenstern, B.; Bauer, G.; Preiser, W.; Doerr, H.W. Stability and inactivation of SARS coronavirus. Med Microbiol Immunol 2005, 194, 1–6, doi: 10.1007/s00430-004-0219-0.

7. Lei, S.; Gao, X.; Sun, Y.; Yu, X.; Zhao, L. Gas chromatography-mass spectrometry method for determination of beta-propiolactone in human inactivated rabies vaccine and its hydrolysis analysis. J Pharm Anal 2018, 8, 373–377, doi: 10.1016/j.jpha.2018.06.003.

8. Hoffmann, M.; Kleine-Weber, H.; Schroeder, S.; Kruger, N.; Herrler, T.; Erichsen, S.; Schiergens, T.S.; Herrler, G.; Wu, N.H.; Nitsche, A., et al. SARS-CoV-2 Cell Entry Depends on ACE2 and TMPRSS2 and Is Blocked by a Clinically Proven Protease Inhibitor. Cell 2020, 181, 271–280 e278, doi: 10.1016/j.cell.2020.02.052.

9. Matsuyama, S.; Nao, N.; Shirato, K.; Kawase, M.; Saito, S.; Takayama, I.; Nagata, N.; Sekizuka, T.; Katoh, H.; Kato, F., et al. Enhanced isolation of SARS-CoV-2 by TMPRSS2-expressing cells. Proc Natl Acad Sci U S A 2020, 117, 7001–7003, doi: 10.1073/pnas.2002589117.

10. Matrosovich, M.; Matrosovich, T.; Garten, W.; Klenk, H.D. New low-viscosity overlay medium for viral plaque assays. Virol J 2006, 3, 63, doi: 10.1186/1743-422X-3-63.

11. Budowsky, E.I.; Friedman, E.A.; Zheleznova, N.V.; Noskov, F.S. Principles of selective inactivation of viral genome. VI. Inactivation of the infectivity of the influenza virus by the action of beta-propiolactone. Vaccine 1991, 9, 398–402, doi: 10.1016/0264-410x(91)90125-p.

12. Budowsky, E.I.; Smirnov Yu, A.; Shenderovich, S.F. Principles of selective inactivation of viral genome. VIII. The influence of beta-propiolactone on immunogenic and protective activities of influenza virus. Vaccine 1993, 11, 343–348, doi: 10.1016/0264-410x(93)90197-6.

13. Chowdhury, P.; Topno, R.; Khan, S.A.; Mahanta, J. Comparison of beta-Propiolactone and Formalin Inactivation on Antigenicity and Immune Response of West Nile Virus. Adv Virol 2015, 2015, 616898, doi: 10.1155/2015/616898.

14. Logrippo, G.A.; Hartman, F.W. Antigenicity of beta-propiolactone-inactivated virus vaccines. J Immunol 1955, 75, 123–128.

15. Gao, Q.; Bao, L.; Mao, H.; Wang, L.; Xu, K.; Yang, M.; Li, Y.; Zhu, L.; Wang, N.; Lv, Z., et al. Rapid development of an inactivated vaccine candidate for SARS-CoV-2. Science 2020, 10.1126/science.abc1932, doi: 10.1126/science.abc1932.

16. Cruz, D.J.; Shin, H.J. Application of a focus formation assay for detection and titration of porcine epidemic diarrhea virus. J Virol Methods 2007, 145, 56–61, doi: 10.1016/j.jviromet.2007.05.012.

17. Muller, C.; Hardt, M.; Schwudke, D.; Neuman, B.W.; Pleschka, S.; Ziebuhr, J. Inhibition of Cytosolic Phospholipase A2alpha Impairs an Early Step of Coronavirus Replication in Cell Culture. J Virol 2018, 92, doi: 10.1128/JVI.01463-17.

18. Ye, C.; Wang, D.; Liu, H.; Ma, H.; Dong, Y.; Yao, M.; Wang, Y.; Zhang, H.; Zhang, L.; Cheng, L., et al. An Improved Enzyme-Linked Focus Formation Assay Revealed Baloxavir Acid as a Potential Antiviral Therapeutic Against Hantavirus Infection. Front Pharmacol 2019, 10, 1203, doi: 10.3389/fphar.2019.01203.

